# Interkingdom Signal Indole Inhibits *Pseudomonas aeruginosa* Persister Cell Waking

**DOI:** 10.1101/674978

**Authors:** Weiwei Zhang, Ryota Yamasaki, Thomas K. Wood

## Abstract

**Aims:** Persister cells are stressed cells that have transient tolerance to antibiotics; these cells undergo no genetic change, but instead, their tolerance is due to reduced metabolism. Unfortunately, little is known about how persisters resuscitate, so we explored the waking of a cells in the presence of the interkingdom signal indole.

**Methods and Results:** To generate a large population of persister cells, we induced the persister phenotype in the opportunistic pathogen *Pseudomonas aeruginosa* by pre-treating cells with carbonyl cyanide *m*-chlorophenylhydrazone to reduce translation by depleting ATP levels, and found, via single cell observations, that proline is sufficient to wake the persister cells. *P. aeruginosa* is often present in the gastrointestinal tract, and indole from commensal bacteria such as *Escherichia coli* has been shown to inhibit *P. aeruginosa* quorum sensing and pathogenicity without influencing growth. Furthermore, indole is not toxic to *P. aeruginosa* persister cells. However, we find here that physiological concentrations of indole inhibit *P. aeruginosa* persister cell resuscitation with an efficiency of higher than 95%. Critically, when contacted with *E. coli* stationary phase cultures, the indole produced by *E. coli* completely inhibits persister cell resuscitation of *P. aeruginosa.*

**Conclusions:** Therefore, *E. col* has devised a method to outcompete its competitors by preventing their resuscitation with indole.

**Significance:** This work provides insight into why indole is produced by commensal bacteria.

## INTRODUCTION

Persister cells become transiently tolerant to antibiotics due to metabolic dormancy, as shown by two groups in the 1940s that discovered the phenotype (Hobby et al., 1942; Bigger, 1944). Unlike resistant cells, which undergo a genetic change and grow in the presence of antibiotics, persister cells do not undergo genetic change (Kwan et al., 2015a; Chowdhury et al., 2016b). Persistence has been found in every strain tested such as *Escherichia coli* (Fisher et al., 2017), *Pseudomonas aeruginosa* (Fisher et al., 2017)*, Staphylococcus aureus* (Fisher et al., 2017) and Archaea (Megaw and Gilmore, 2017). Also, both antibiotic and nutrient stress creates persister cells (Bernier et al., 2013; Maisonneuve and Gerdes, 2014; Martins et al., 2018). Since virtually all cells experience nutrient stress (Song and Wood, 2018) and since the viable but not culturable state appears to be the same as the persister state (Kim et al., 2018b), persistence may be a universal phenotype that is the key to microbial survival.

Stress likely activates toxins of toxin/antitoxin (TA) systems to render the metabolic dormancy of the persister state since deletion of several toxins reduces persistence (Harrison et al., 2009; Dörr et al., 2010; Kim and Wood, 2010; Luidalepp et al., 2011) and production of toxins, from TA systems (Hong et al., 2012) or toxins unrelated to TA systems (Chowdhury et al., 2016a), increases persistence dramatically. Persisters wake over a range of times from instantaneously to several hours as a function of their ribosome content (Kim et al., 2018a). However, how nutrients are perceived and how persisters resuscitate is not understood well.

The low numbers of persister cells in bacterial populations (about 1% in biofilms and stationary-phage cultures (Lewis, 2007; Lewis, 2008) hinders our understanding of these cells. We have developed methods to increase the persister cell population to nearly 100% by pre-treating with acid or hydrogen peroxide (Hong et al., 2012) as well as by halting RNA transcription via rifampicin, by stopping translation via tetracycline, and by ceasing ATP production via carbonyl cyanide *m*-chlorophenylhydrazone (CCCP) (Kwan et al., 2013). The persister cells generated by rifampicin pretreatment have been validated via eight different assays (multi-drug tolerance, immediate change from persister to non-persister in the presence of nutrients, two lines of evidence for dormancy, no change in the minimum inhibitory concentration compared to exponential cells, no resistance phenotype, similar morphology to ampicillin-induced persisters and similar resuscitation as ampicillin-induced persisters) (Kim et al., 2018a). Our chemical methods have been validated by six independent groups in recent months (Grassi et al., 2017; Cui et al., 2018; Narayanaswamy et al., 2018; Sulaiman et al., 2018; Tkhilaishvili et al., 2018; Pu et al., 2019) and have been shown to be effective not only for *E. coli* but also for *P. aeruginosa* and *S. aureus.*

Indole is an interkingdom signal since it is produced by commensal bacteria and once it is detected by epithelial cells, they use it to tighten their cell junctions to reduce infection (Bansal et al., 2010). In addition, indole produced by rhizosphere bacteria stimulates *Arabidopsis thaliana* plant growth (Blom et al., 2011), and indole has long been recognized for affecting insect behavior (Dethier, 1947). As an interspecies signal, indole produced by *E. coli* reduces quorum-sensing-related virulence factors in *P. aeruginosa* (Lee et al., 2009), and indole increases the competiveness of *E. coli* in dual cultures with *P. aeruginosa* (Chu et al., 2012). This is significant since the opportunistic pathogen *P. aeruginosa* is better known as a respiratory and wound pathogen; however, *P. aeruginosa* would experience indole since it is present in the gastrointestinal (GI) tract in up to 12% of the normal population (Bodey et al., 1983), and its presence in the GI tract of critically-ill surgical patients results in a nearly three-fold increase in mortality (Marshall et al., 1993).

In regard to persister cells, indole (Hu et al., 2015; Kwan et al., 2015b) and substituted indoles (Lee et al., 2016) have been shown to kill *E. coli* persister cells; however, they do not kill *P. aeruginosa* persisters (Lee et al., 2016). Critically, the effect of indole on persister cell waking has not been investigated previously. Here, we find that although indole does not affect *E. coli* persister cell waking (i.e., indole does not affect the cell that produces it), indole nearly completely prevents *P. aeruginosa* persister cell waking without affecting its growth.

## METHODS

### Bacteria and culture conditions

Strains (**Table 1**) were grown routinely in LB medium (Sambrook, 1989) at 37 °C. M9 minimal medium was supplemented with different amino acids (Rodriguez., 1983). M9-casamino acid tryptophan was prepared as described previously (Li, 2013) and used for indole production. CCCP was dissolved in DMSO at stock a concentration of 50 mg/mL. Indole was dissolved in dimethyl sulfoxide at a stock concentration of 1 M.

**Table 1.**
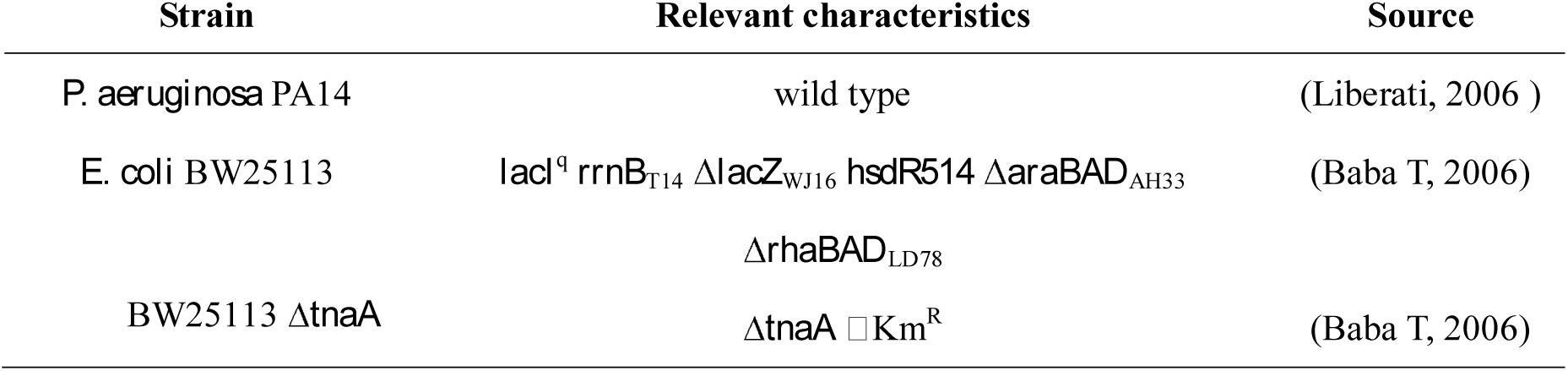
Strains used in this study.

### Persister cells

PA14 persister cells were generated by stopping ATP production with CCCP (Kwan et al., 2013) and by lysing non-dormant cells with ciprofloxacin. One milliliter of overnight PA14 culture was diluted in 5 mL of fresh LB with 24 µL of CCCP stock solution (final concentration 200 µg/mL) and incubated for 3 h. Bacteria were washed with 0.85% NaCl (3500 × g for 10 min), resuspended in LB supplemented with ciprofloxacin (5 µg/mL) for 3 h to lyse non-persister cells, then washed twice with 0.85% NaCl.

*E. coli* BW25113 (hereafter BW25113) persister cells were generated by stopping transcription with rifampicin (Kwan et al., 2013). Overnight cultures of BW25113 were inoculated into 25 mL of LB and grown to a turbidity of 0.8. Rifampicin was added (100 µg/mL) for 30 min, cells were centrifuged, and resuspended in LB supplemented with (100 µg/mL ampicillin) and incubated for 3 hours.

### PA14 resuscitation with amino acids

To observe resuscitation of *P. aeruginosa* PA14 (hereafter PA14) persister cells with amino acids as the sole carbon source, amino acids were initially divided into four combinations of five each in M9 minimal medium: combination #1: Arg, Ser, Cys, Ala and Phe,, combination #2: His, Thr, Gly, Val, and Tyr, combination #3:Lys, Asn, Pro, Ile, and Trp, and combination #4:Asp, Glu, Gln, Leu, and Met. To observe the resuscitation of persister cells on single amino acids, the 15 amino acids from combinations 2, 3, and 4 were tested separately. PA14 persister cells were diluted 5000-fold, and 50 μL was spread onto M9 plates containing each kind of individual amino acid.

### Single cell resuscitation

To observe the resuscitation of single cells, agarose gel pads were used (Kim et al., 2018a). Briefly, low melting agarose (1.5 %) (Nusieve GTG agarose BMB # 50081) was added to 11 mL H_2_O and melted by microwaving (30 sec at 1,500 Watts), and 1.25 mL 10×salt solution (0.125 mL MgSO_4_ and 0.125 mL CaCl_2_) with proline (0.164 mg/mL) was added after cooling. Single cells were observed using light microscope (Zeiss Axioscope.A1, bl_ph channel at 1,000 ms exposure) maintained at 37 °C.

### *P. aeruginosa* resuscitation in the presence of *E. coli*

PA14 persister cells (1 mL) were added to 25 mL cultures of BW25113 grown in M9 casamino acids tryptophan medium for 24 hrs. The mixtures were incubated at 37 °C with shaking and samples taken after 5 min, 30 min, 60 min and 120 min. Cells were diluted 10^4^- to 10^6^-fold, washed twice with 0.85 % NaCl, and plated on LB plates. PA14 and BW25113 were enumerated based on color discrimination and the antibiotic resistance (PA14 has natural resistance to ampicillin).

## RESULTS

### CCCP-induces PA14 persister formation

In order to study PA14 resuscitation, we first optimized conditions for PA14 persister cell formation, using methods we developed for *E. coli* to reduce ATP and thereby induce the dormant state (Kwan et al., 2013). We found that 200 µg/mL pretreatment with CCCP for 3 h induced PA14 persister cell formation with an efficiency of 98.6% as evidenced by their ciprofloxacin tolerance (**Fig. S1**). To verify that these cells are *bona fide* persister cells, we completed three additional assays and found (i) re-grown persister cells have the same sensitivity to the antibiotic (5 μg/mL ciprofloxacin, **Fig. S2**), (ii) CCCP-generated persisters show the same minimal inhibitory concentration as the wild type PA14 (**Fig. S3**), and (iii) 99.3% of the PA14 persister cells obtained from CCCP-pretreatment do not resuscitate in 10 h on agarose pads lacking nutrients (1 cell out of 228 resuscitate) (**Fig. 1A**) whereas 100% of exponentially-grown PA14 cells resuscitate on gel pads without any carbon source (**Fig. 1B**). Therefore, CCCP-pretreatment induces PA14 into the persister state with high efficiency.

**Fig. 1.**
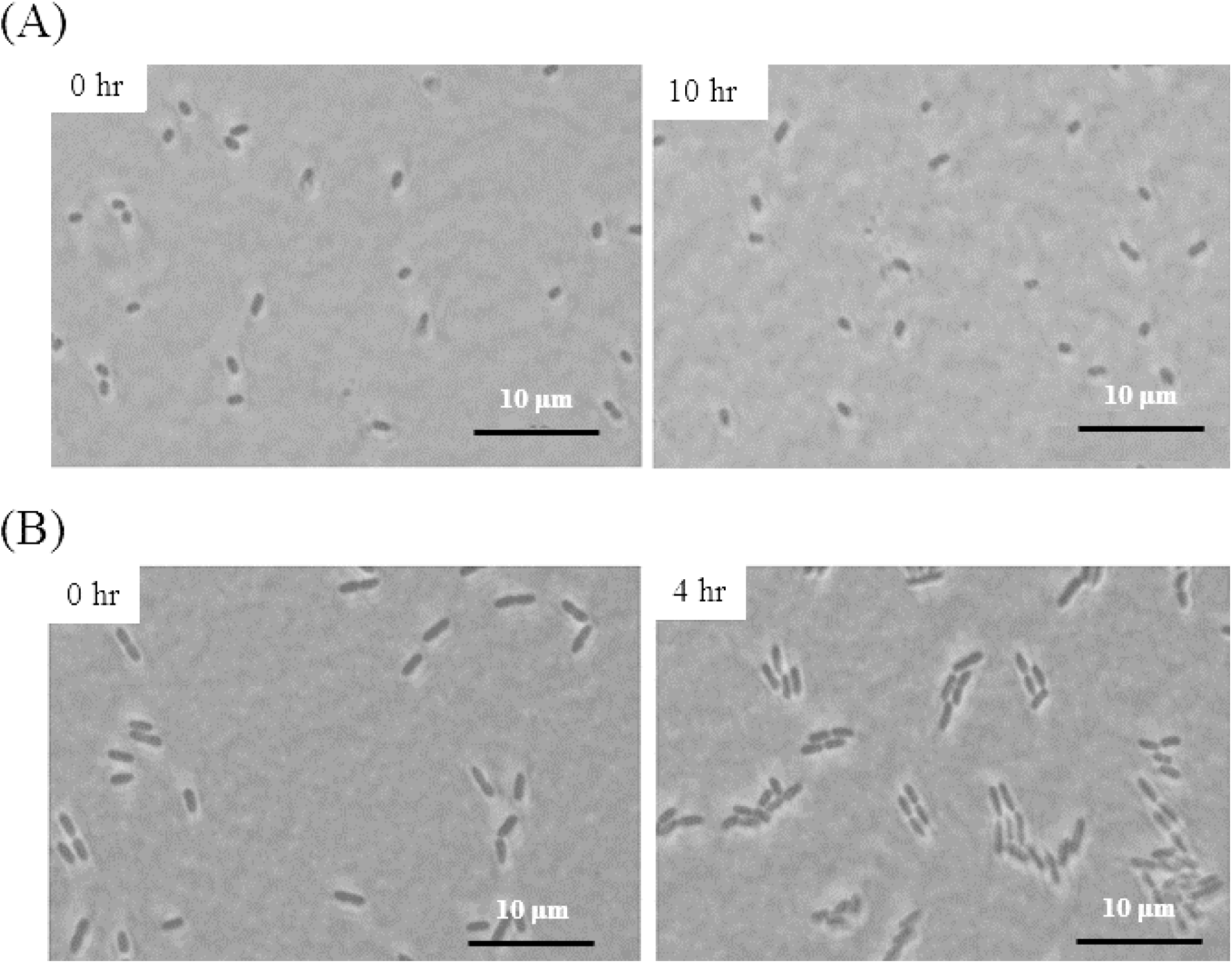
PA14 persister cell waking in the absence of a carbon source. (**A**) Resuscitation of single PA14 persister cells on M9 medium gel pads that lack a carbon source after 10 h. (**B**) As a positive control, exponentially-grown PA14 cells were tested for resuscitation after 4 h in the absence of a carbon source. Scale bar indicates 10 µm. One of two independent cultures shown.

### PA14 persister cells resuscitate on proline

Having established how to generate PA14 persister cells, we investigated whether a single amino acid wakes PA14 persisters since alanine revives *Bacillus subtilis* spores (Mutlu et al., 2018); our hypothesis is that dormant PA14 persister cells wake like dormant spores.

To rapidly determine which amino acid wakes PA14 persisters, for the initial assay, four groups of five amino acids as the carbon and nitrogen source were screened : combination #1: Arg, Ser, Cys, Ala and Phe,, combination #2: His, Thr, Gly, Val, and Tyr, combination #3:Lys, Asn, Pro, Ile, and Trp, and combination #4:Asp, Glu, Gln, Leu, and Met. Using an agar plate assay in which persister cells that wake first form larger colonies, we found that PA14 persister cells do not revive with amino acid combination #1 but revive well with combination #3 and revive to a lesser extent with combinations #2 and #4 (**Fig. S4** and **Table S1**). We then proceeded to separately test each of the 15 amino acids from combinations #2, #3 and #4, and found PA14 persister cells wake fastest with proline as the sole carbon source (**Fig. 2**). Using observations of single cells waking on proline, we found that 88% of the PA14 persisters revive in 10 h on M9 proline agar plates (**Fig. 3**).

**Fig. 2.**
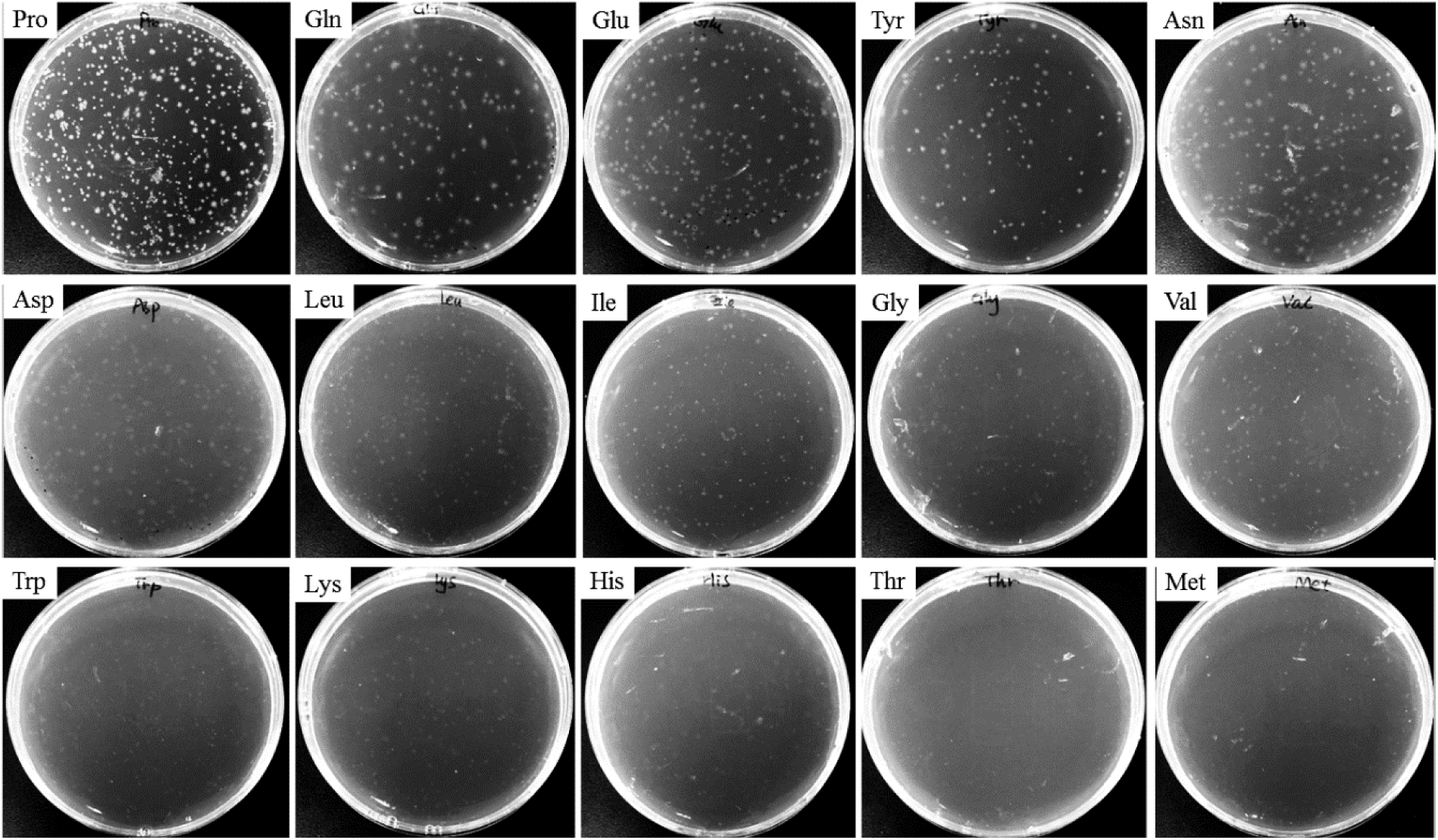
PA14 persister cell waking on single amino acids. Resuscitation of PA14 persister cells after 48 h at 37°C on M9 agar medium with 15 individual amino acids.

**Fig. 3.**
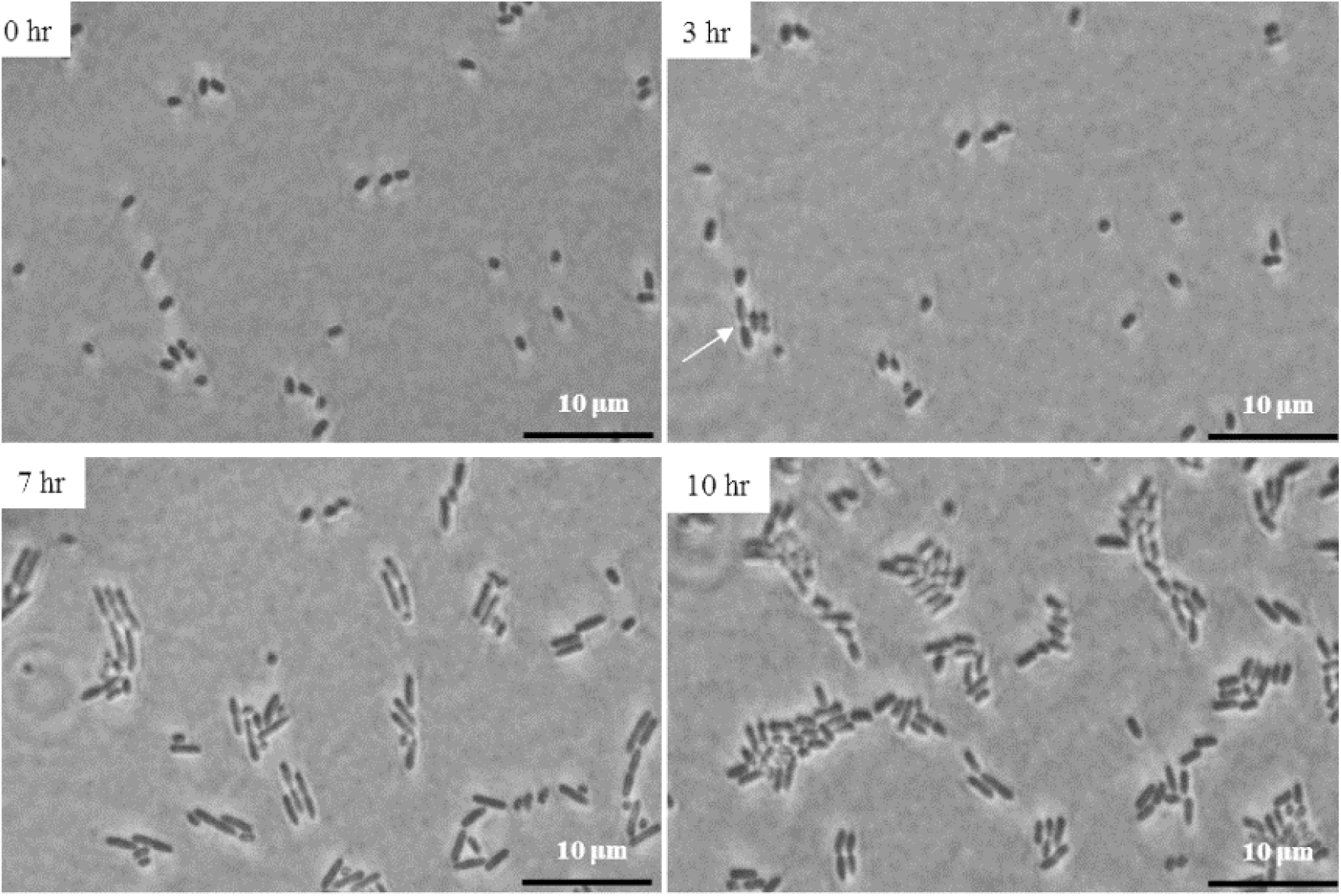
PA14 persister cell waking on proline. Resuscitation of single PA14 persister cells after 10 h at 37°C on M9 proline agarose pads. The white indicates one of the first cells to resuscitate. Scale bar indicates 10 µm. One of two independent cultures shown.

### Indole inhibits PA14 persister cell waking

Since indole inhibits the virulence of *P. aeruginosa* (Lee et al., 2016), we hypothesized that indole may affect the resuscitation of PA14 persister cells. We found that when 2 mM indole is present, it inhibits the waking of more than 95% of the PA14 persister cells (**Fig. S5**). Corroborating this result with single cells, we found 99% of PA14 persister cells were inhibited after 10 h on M9 proline plates containing 2 mM indole (**Fig. 4**). Critically, 2 mM indole was not toxic to PA14 persister cells (**Fig. S6**) (the specific growth rate was reduced only by 5.8%) so the inhibition of PA14 persister cell resuscitation was not due to growth inhibition. Indole was also not toxic to BW25113 persister cells and unlike PA14, indole had no inhibitory effect on the wake up of BW25113 persister cells (**Fig. S7**).

**Fig. 4.**
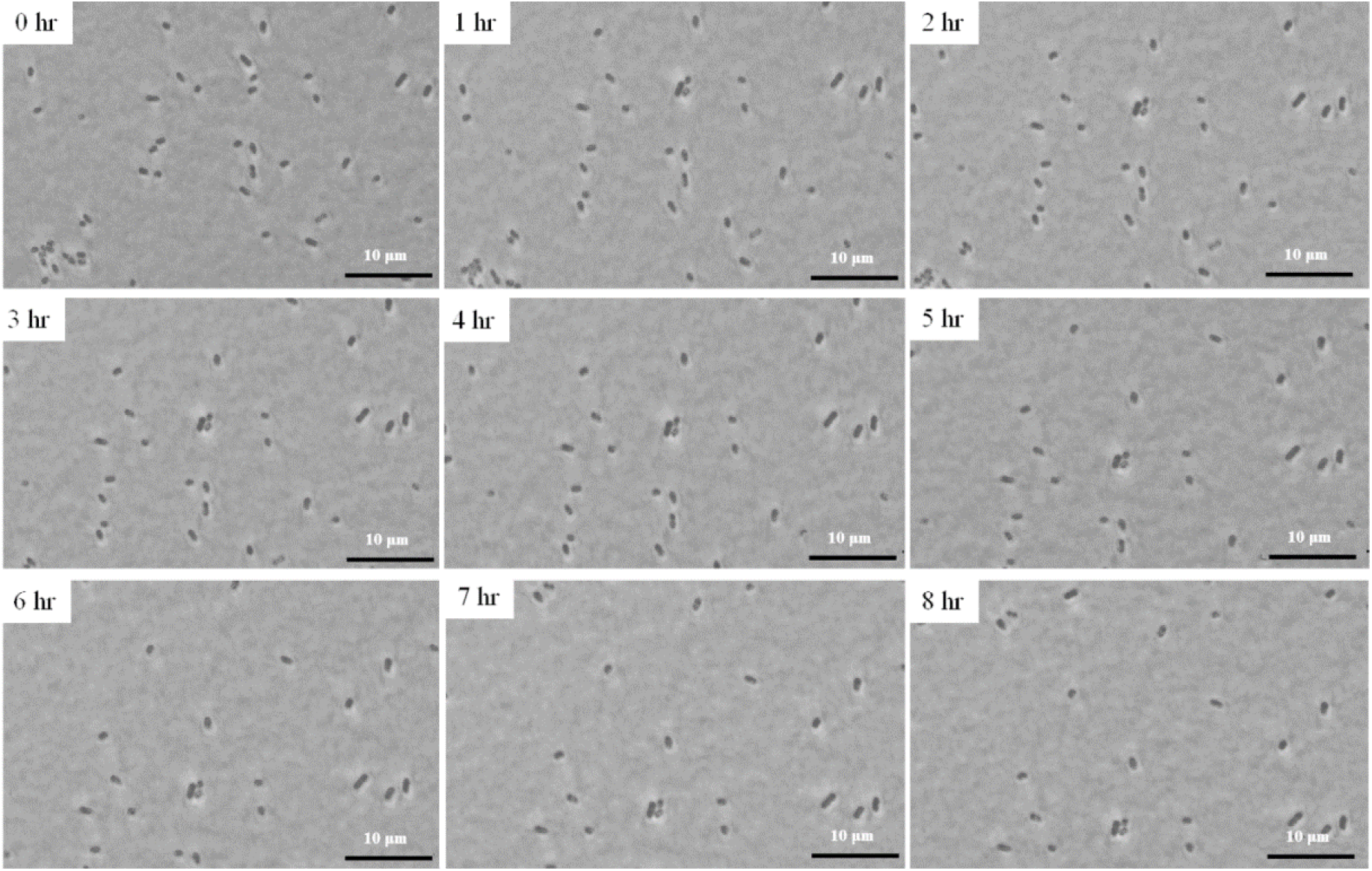
Indole inhibits PA14 persister cell waking on proline. Resuscitation of single PA14 persister cells after 10 h on M9 proline + 2 mM indole agarose gel pads. Scale bar indicates 10 µm. One of two independent cultures shown.

### BW25113 producing indole outcompetes PA14

BW25113 grown in M9 casamino acids tryptophan medium produced 4.6 mM indole in the culture supernatant after 24 h, while no indole was produced by the *E. coli* BW25113△*tnaA* since it lacks tryptophanase (**Table S2**). When PA14 persister cells were added to the BW25113 cells grown in this medium, no PA14 persister cells resuscitated. Without indole, 97.1%, 93.4%, 98.8%, and 98.7% of the PA14 persister cells resuscitated in the *E.* coli △*tnaA* culture after 5, 30, 60, and 120 min (**Fig. 5**). Hence, indole produced by *E. coli* prevents *P. aeruginosa* persister cells from resuscitating.

**Fig. 5.**
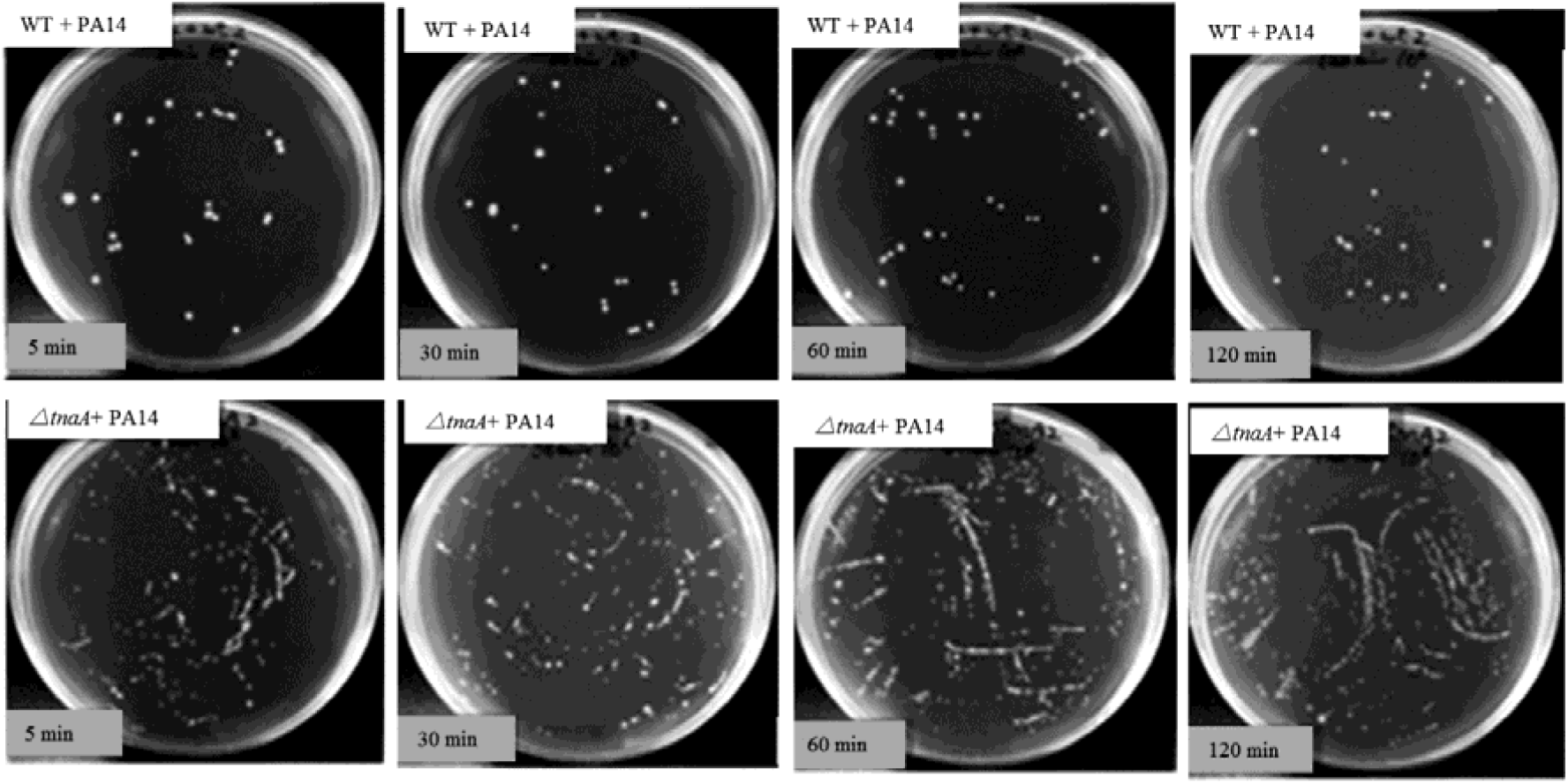
Indole gives *E. coli* a competitive advantage over PA14. PA14 persister cells were contacted with *E. coli* BW25113 wild type (**A**) and with *E. coli* BW25113 Δ*tnaA* (**B**) M9 casamino acids tryptophan medium cultures for 5, 30, 60 and 120 min, then samples were added to LB plates that were incubated at 37 °C for 16 hours.

## DISCUSSION

By preventing protein production via a CCCP pretreatment followed by contacting with ciprofloxacin to lyse any non-dormant cells, we converted *P. aeruginosa* cultures into nearly homogeneous persister cell populations; hence, we converted the rare persister phenotype (less than 1%) into the dominant phenotype. By using this approach, we were able to determine that *P. aeruginosa* persister cells can be resuscitated by the amino acid proline. Building on this discovery, we were able to determine a new physiological role for the interkingdom signal indole: *E. coli* cells in the GI tract produce indole to not only tighten human epithelial cell junctions (Bansal et al., 2010) but indole is also produced by commensal *E. coli* to prevent *P. aeruginosa* persister cells from resuscitating.

This discovery of indole inhibiting *P. aeruginosa* persister cell waking is physiologically relevant since *P. aeruginosa* is found in the GI tract (Bodey et al., 1983; Marshall et al., 1993), and cells in the GI tract experience nutritional stress between meals. This stress likely leads to the formation of *P. aeruginosa* persister cells in the GI tract. Therefore, our results suggest that indole is produced by *E. coli* as a tool to prevent *P. aeruginosa* cells from waking so that *E. coli* cells may outcompete *P. aeruginosa* cells for limited nutrients by resuscitating first.

## ACKNOWLEDGEMENTS

This work was supported by funds derived from the Biotechnology Endowed Professorship at the Pennsylvania State University. This work also was supported by National Natural Science Foundation of China (41676141), the China Scholarship Council (1712290032) and the K.C. Wong Magna Fund at Ningbo University, China. The authors confirm there are no conflicts of interest.

## Supporting Information

**Table S1.** PA14 persister resuscitation on M9 minimal agar plates with groups of five amino acids. Colonies were counted after 24 and 48 h.

| **Amino acid**<br>**combination** | **Colony number**<br>**(24 h)** | **Colony number**<br>**(48 h)** |
| --- | --- | --- |
| No amino acid | 0 | 0 |
| #1 | 0 | 0 |
| #2 | 30 ± 16 | 49 ± 25 |
| #3 | 44 ± 17 | 74 ± 16 |
| #4 | 18 ± 1 | 53 ± 21 |

**Table S2.** Indole produced by *E. coli* BW25113 in in M9 casamino acids tryptophan medium. Cells were cultured for 24 h, and the turbidity at 600 nm is indicated. Two independent cultures were used (labeled as #1 and #2).

| **Samples** | **Turbidity** | **Indole in supernatant**<br>**($\mu\text{M}$)** |
| --- | --- | --- |
| **WT #1** | 3.6 | 501 ±40 |
| **WT #2** | 3.3 | 448 ± 5 |
| **$\Delta tnaA$ #1** | 3.7 | 0 |
| **$\Delta tnaA$ #2** | 3.5 | 0 |

**Fig. S1.**
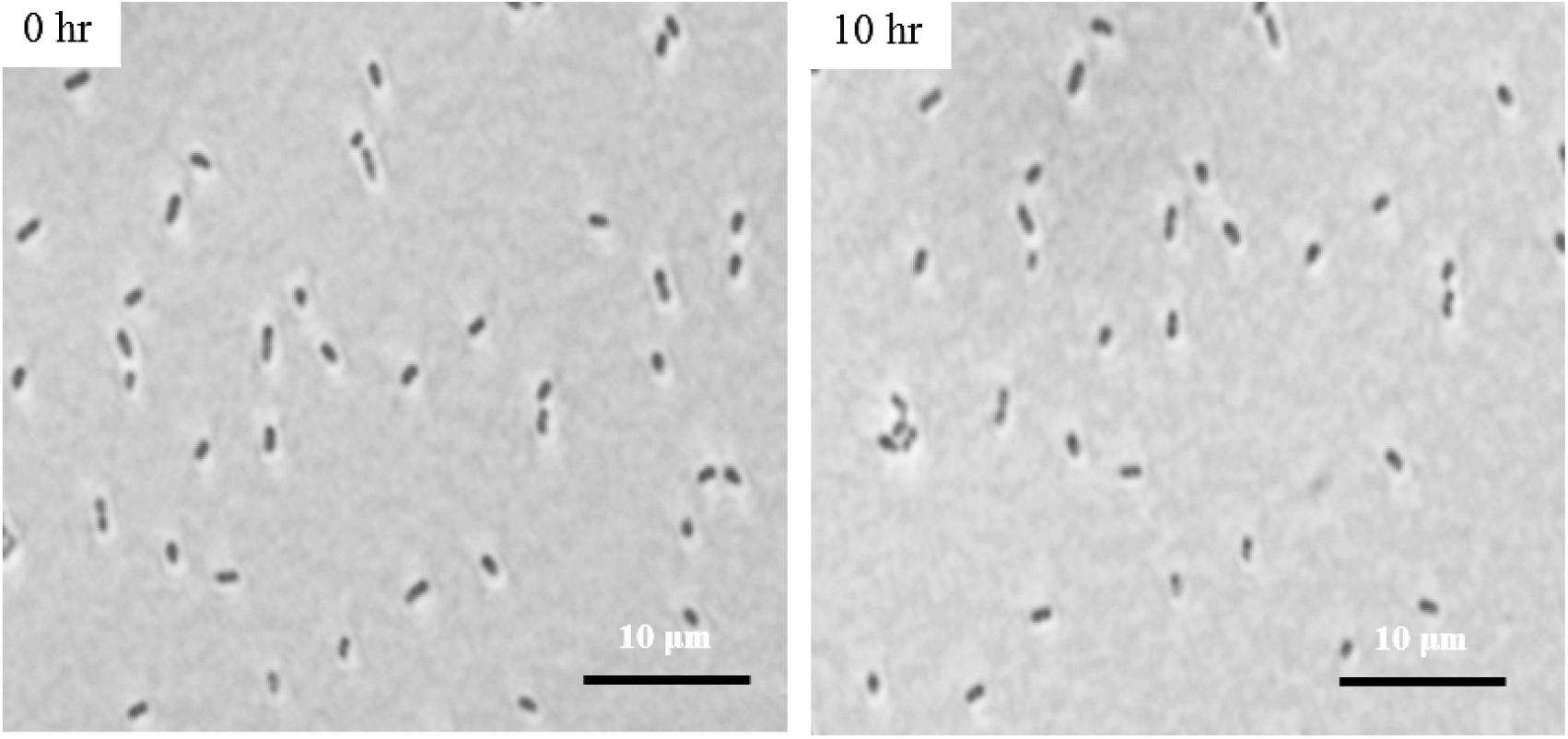
CCCP-generated *P. aeruginosa* PA14 persister cells are tolerant to ciprofloxacin. Persister cells on the M9 gel pads (no amino acids) with ciprofloxacin do not lyse in 10 h. For the two independent cultures, 1 out of 122 cells woke (i.e., elongated and died due to the antibiotic) and 1 out of 191 cells woke. One representative sample shown.

**Fig. S2.**
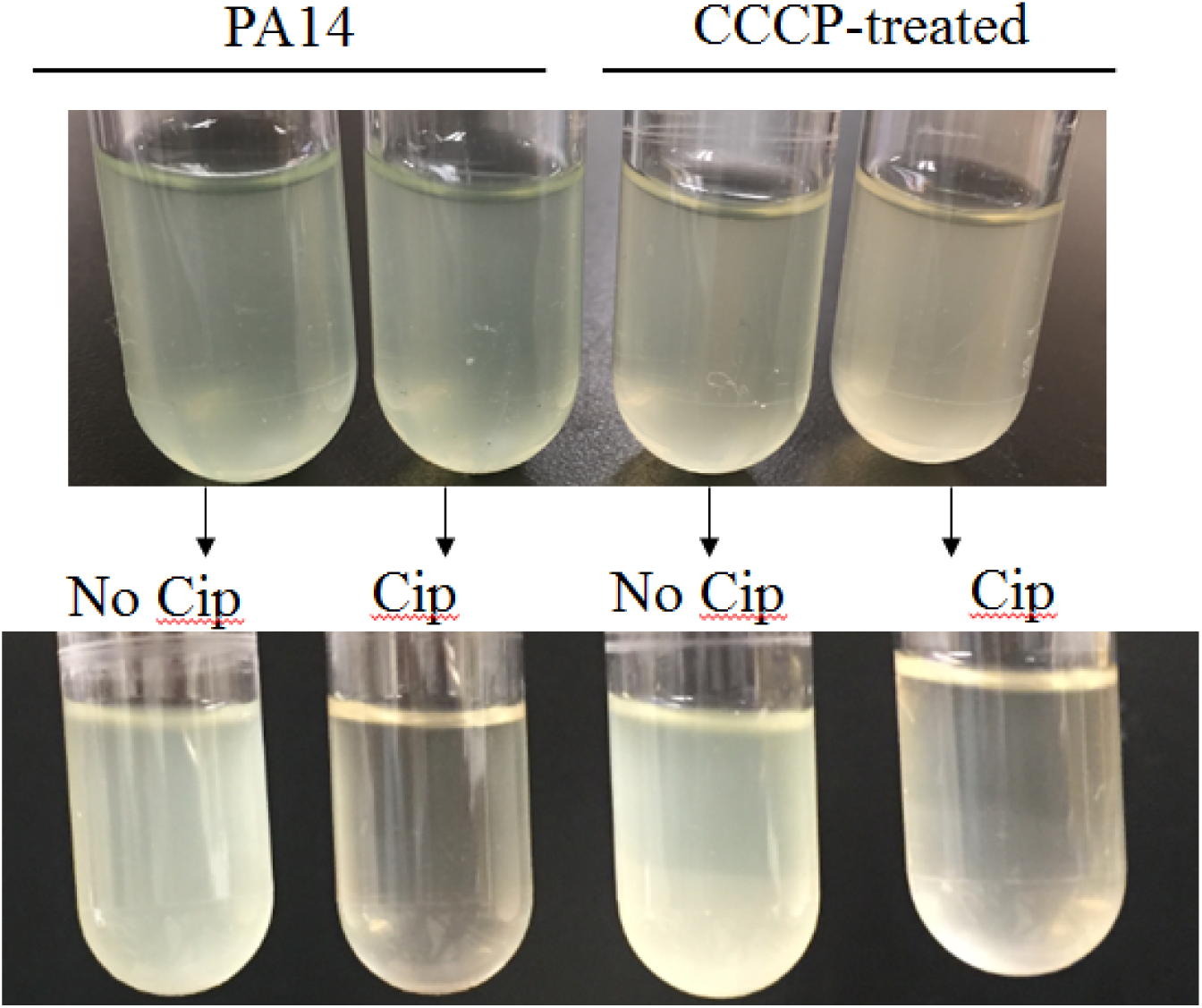
CCCP-generated *P. aeruginosa* PA14 persister cells have the same antibiotic tolerance as wild-type cells. After re-growth in LB medium, CCCP-generated persister cells are lysed by 5 μg/mL ciprofloxacin in the same manner as wild-type PA14 cells.

**Fig. S3.**
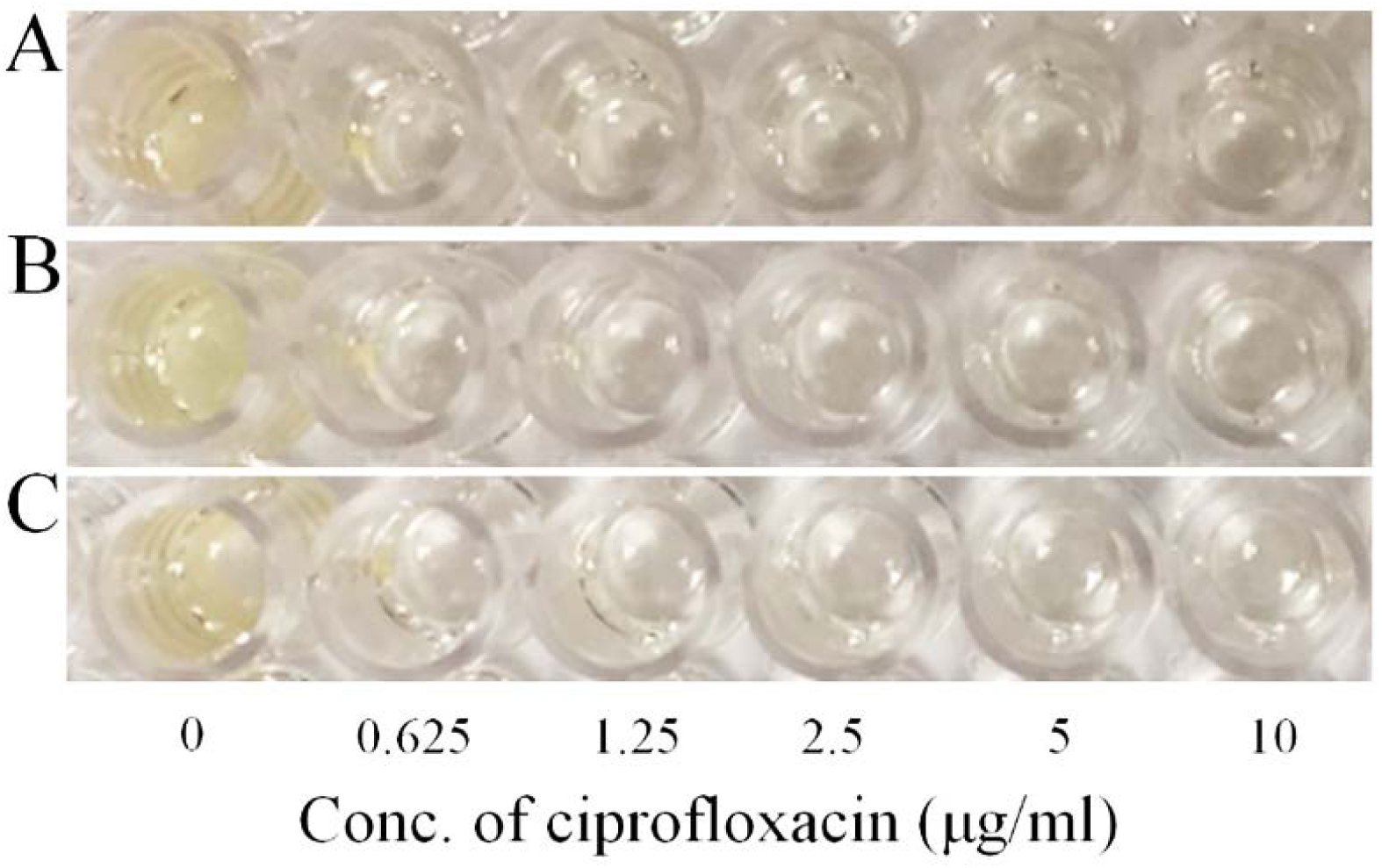
CCCP-generated *P. aeruginosa* PA14 persister cells have the same minimum inhibitory concentration as wild-type cells. Overnight cultures of PA14 at turbidity 4.0 (**A**), CCCP-induced persister cells at turbidity 3.6 (without ciprofloxacin treatment) (**B**), and re-grown CCCP-induced persisters at turbidy 2.5(**C**) were diluted into fresh LB at a concentration of 10^4^ cells/mL. Cells (196 μL) were placed in 96-well plates, and ciprofloxacin was added in 4 μL. The final concentrations of ciprofloxacin were 0, 0.625, 1.25, 2.5, 5, and 10 µg/ml as indicated. The 96-well plates were incubated at 37 °C for overnight, and the concentration at which cells did not grow was used as the MIC.

**Fig. S4.**
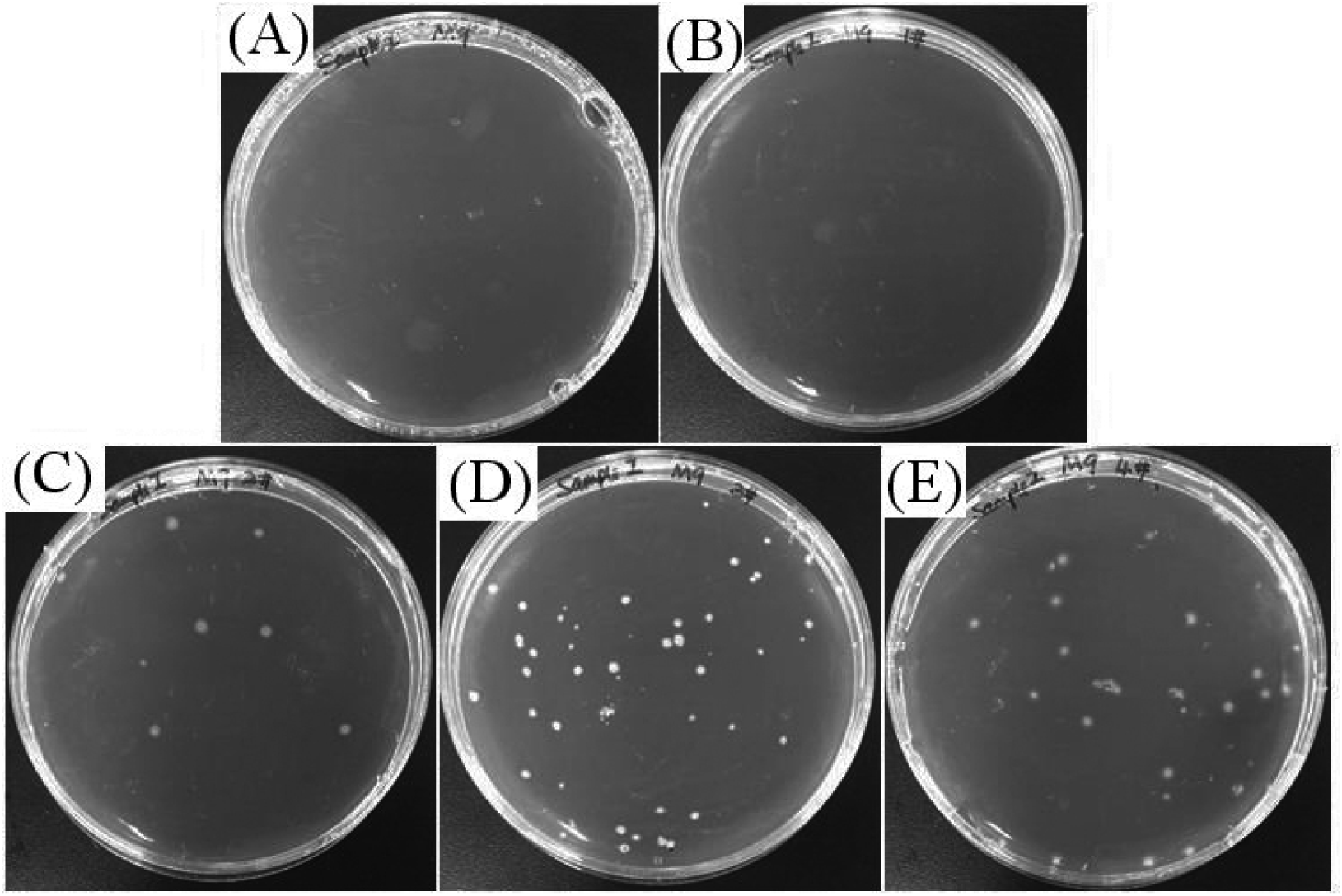
PA14 persister cell resuscitation on groups of five amino acids. PA14 persister cells were grown for 48 h on M9 agar plate supplemented with different combinations of five amino acids as the sole carbon source: (**A**) M9 minimal medium plates without amino acids; (**B**) M9 with combination #1; (**C**) M9 with combination #2; (**D**) M9 with combination #3; and (**E**) M9 with combination #4. Specific cell numbers are shown in **Table S1.**

**Fig. S5.**
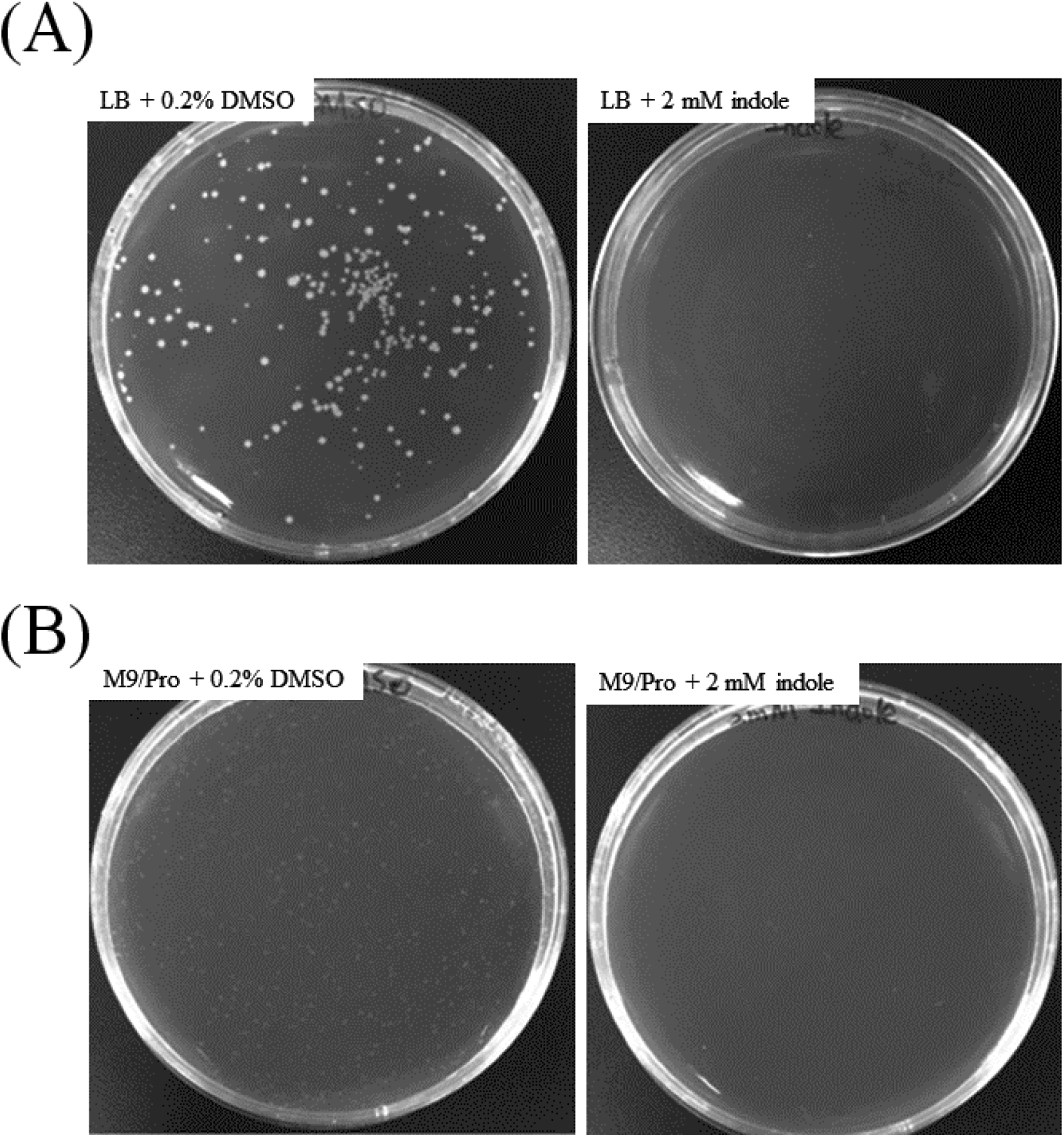
Indole inhibits PA14 resuscitation. PA14 resuscitation after one day on LB + 0.2% DMSO and LB + 2 mM indole (**A**), and on M9 proline + 0.2% DMSO and M9 proline + 2 mM indole (**B**).

**Fig. S6.**
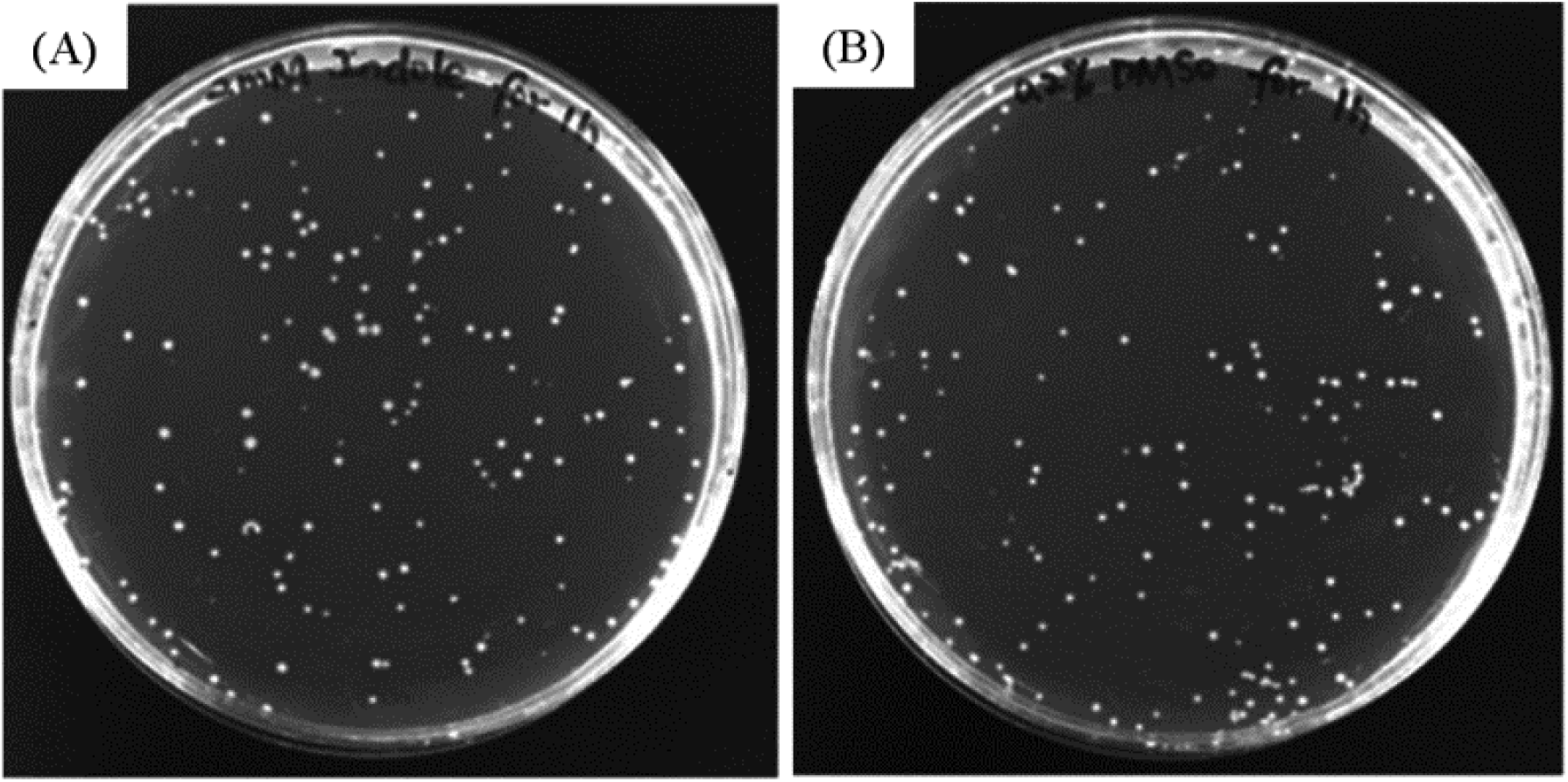
Indole is not toxic to PA14 persister cells. PA14 persister cells were treated for 1 h with 2 mM indole (**A**) and 0.2% DMSO (**B**), diluted, and plated (50 μL) on LB plates to check for cell viability.

**Fig. S7.**
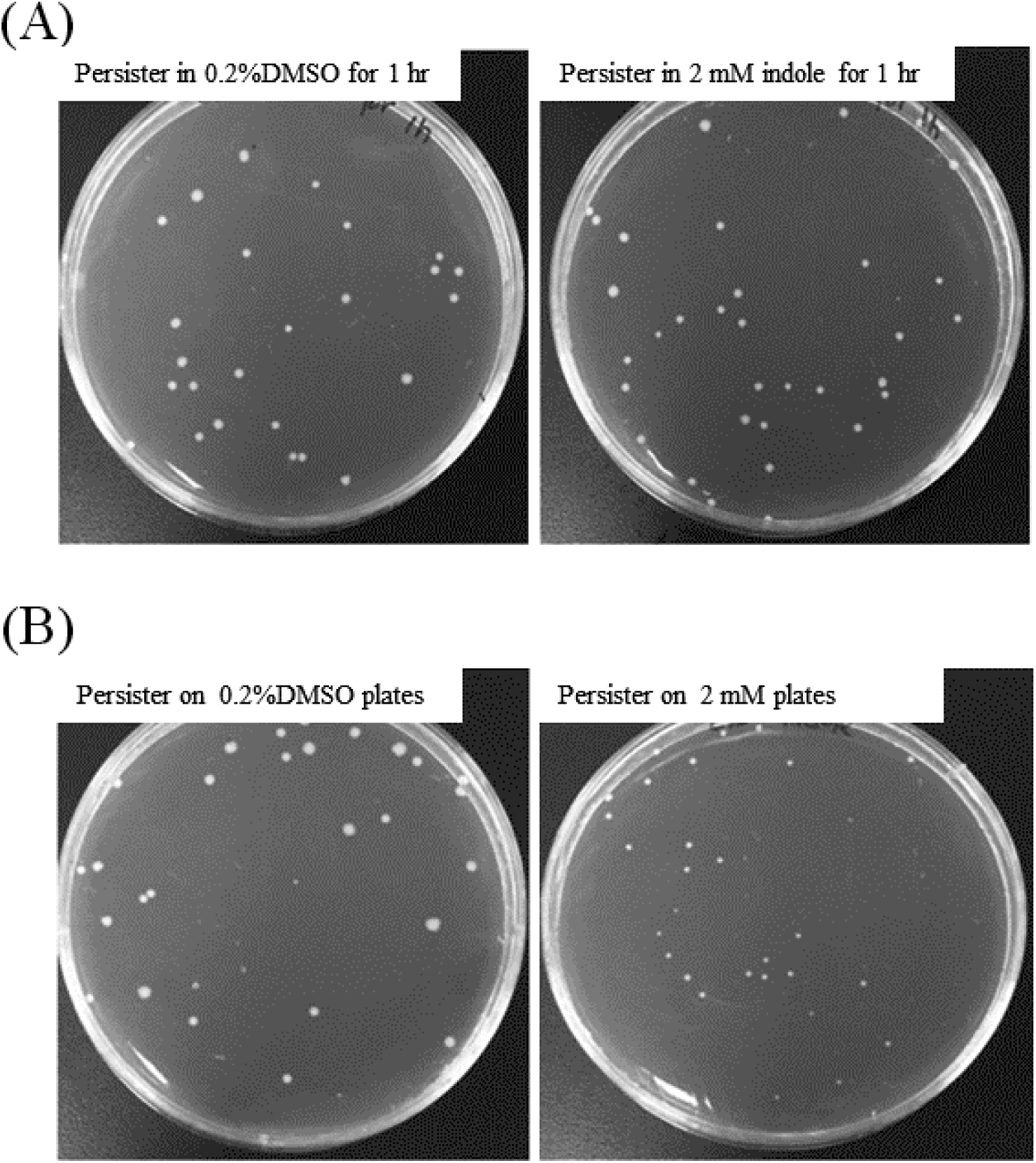
Indole is not toxic to *E. coli* persister cells. (**A**) *E. coli* persister cells were treated with 2 mM indole or 0.2% DMSO for 1 h, diluted, and plated (50 μL) on LB plates. (**B**) Resuscitation of *E. coli* persister cells on LB + 0.2%DMSO and LB + 2 mM indole plates.

